# Prevalence of Bourbon and Heartland viruses in field collected ticks at an environmental field station in St. Louis County, Missouri, USA

**DOI:** 10.1101/2022.03.23.485543

**Authors:** Ishmael D Aziati, Derek McFarland, Avan Antia, Astha Joshi, Anahi Aviles-Gamboa, Preston Lee, Houda Harastani, David Wang, Solny A. Adalsteinsson, Adrianus C. M. Boon

## Abstract

Heartland and Bourbon viruses are pathogenic tick-borne viruses putatively transmitted by *Amblyomma americanum*, an abundant tick species in Missouri. To assess the prevalence of these viruses in ticks, we collected 2778 ticks from 8 sampling sites at Tyson Research Center, an environmental field station within St. Louis County and close to the City of St. Louis, from May - July in 2019 and 2021. Ticks were pooled according to life stage and sex, grouped by year and sampling site to create 355 pools and screened by RT-qPCR for Bourbon and Heartland viruses. Overall, 14 (3.9%) and 27 (7.6%) of the pools were positive for Bourbon virus and Heartland virus respectively. In 2019, 11 and 23 pools were positive for Bourbon and Heartland viruses respectively. These positives pools were of males, females and nymphs. In 2021, there were 4 virus positive pools out of which 3 were positive for both viruses and were comprised of females and nymphs. Five out of the 8 sampling sites were positive for at least one virus. This included a site that was positive for both viruses in both years. Detection of these viruses in an area close to a relatively large metropolis presents a greater public health threat than previously thought.

## INTRODUCTION

Emerging infectious diseases (EIDs) have become an increasing global concern in the twenty-first century; their emergence and spread pose a significant threat to global health and economies [1]. From an extensive review of the literature for EID events between 1940 and 2004, the majority (60.3%) were caused by zoonotic pathogens. Remarkably, the second (22.8%) most important category was infections caused by vector-borne diseases [2]. Vector-borne infections are an important cause of morbidity and mortality with mosquitoes being responsible for most of the cases worldwide. In the United States (US) however, ticks are the leading disease vector accounting for over 90% of the annual vector-borne cases with Lyme disease being the most commonly reported [3–5]. Ticks are blood-sucking arthropods that are competent vectors of a wide range of vertebrate pathogens including viruses [6, 7]. Substantial geographic expansions of tick populations have been revealed by tick surveys throughout the US, and tick species like the lone star tick (*Amblyomma americanum*) have expanded their range in the US in recent decades [8, 9]. The lone star tick is responsible for the continuous emergence and spread of tick-borne diseases like ehrlichiosis, tularemia and STARI [4]. Aside from these, field isolations and laboratory studies have implicated *A. americanum* tick as the putative vector of Heartland virus (HRTV) and Bourbon virus (BRBV), two tick-borne viruses discovered in the US in the last decade [10–15]. HRTV and BRBV were first isolated from febrile patients in Missouri (MO) and Kansas (KS) respectively, who reported histories of tick bites [16, 17]. Since the discovery of these new tick-borne viruses, about 50 HRTV and 5 BRBV infections have been confirmed in human cases of febrile illness with sometimes fatal outcome across the US [11, 18, 19]. Similarly, these viruses have been detected in field collected *A. americanum* ticks, along with serological evidence of infection in wildlife in the Midwest and Eastern US states [13–15, 18, 20–22].

BRBV is a segmented RNA virus that belongs to the family *Orthomyxoviridae* and the genus *Thogotoviru*s. Members of this genus have been shown to be distributed worldwide but not until recently in the US [12, 14, 16, 18]. BRBV is the first *Thogotovirus* discovered in North America that is known to infect humans. Three other members of this genus, *Thogotovirus*, Oz virus and Dhori virus, which have not been found in the US, have been associated with human disease [23–25]. HRTV is also a segmented RNA virus that belongs to the family *Bunyaviridae:* genus *Phlebovirus* [26]. It is genetically closely related to Dabie bandavirus, a virus that was first identified in China in 2009 and later reported in Japan and South Korea [11, 27].

The prevalence of BRBV and HRTV in field collected ticks has only been reported in the rural northwestern part of Missouri, with no peer-reviewed published reports from the other parts of the state. As part of a broader study aimed at an integrated vector-animal-human surveillance for known and novel tick-borne viruses, we tested the prevalence of BRBV and HRTV in field collected ticks from Tyson Research Center in Missouri an environmental field station that is known to have abundant host-seeking *A. americanum* ticks as well as co-mingling of vectors and different animal hosts [28]. We determined the prevalence of HRTV and BRBV by RT-qPCR in *A. americanum* sampled in 2019 and 2021 and report for the first time the surveillance and detection of these viruses in an area close to a relatively large metropolitan area. The infection rates per 1000 ticks varied among collection sites, life stages, sex, and between collection years.

## MATERIALS AND METHODS

### Study Area

Tyson Research Center (TRC) is the ~800-ha environmental field station of Washington University in St. Louis, situated ~20 km southwest of the city of St. Louis, Missouri, US within the adjacent St. Louis County (38°31ŪN, 90°33ūW). With the exception of an interstate to its southern edge, TRC is surrounded by protected lands that are popular with outdoor recreationists. TRC is located in the northeastern edge of the Ozark ecoregion and is ~85% forested, consisting of mainly deciduous oak-hickory forest on steep slopes and ridges. South-facing slopes tend to be dominated by chinquapin oak (*Quercus muehlenbergii*) and eastern red cedar (*Juniperus virginiaiia*); protected slopes by flowering dogwood (*Cornus florida*), white oak (*Q. alba*), and black oak (*Q. velutina*); and bottomlands by slippery elm (*Ulmus rubra*) and American sycamore (*Plantanus occidentalis*) [29]. Forest understory contains areas of woody shrubs including spicebush (*Lindera benzoin*), common buckthorn (*Frangula caroliniana*), and pawpaw (*Asimina triloba*). In all habitat types there are significant and patchy areas of Amur honeysuckle (*Lonicera maackii*) invasion, some of which are under ongoing management. TRC also maintains open old field habitat in its central valley and has ~24 ha of limestone/dolomite glades that are heavily invaded by red cedar. Prior to its acquisition by Washington University in St. Louis, TRC land has been used for: mining (at Mincke Quarry), logging, grazing livestock, habitation of a small village (now only ruins/building foundations remain in Mincke Valley), and construction of dozens of bunkers (which remain along the central valley) for munitions storage by the US Military. *Amblyomma americanum* is the most commonly-encountered tick species at TRC; *Dermacentor variabilis* and *Ixodes scapularis* are also present [28]

### Tick Collection and Identification

Ticks were collected between May and July in 2019 and 2021 from eight different locations within TRC. These included Bunker 51, Bunker 37, Bat Road, Library Road, North Gate, Mincke Quarry, Mincke Valley and Plot 7/8 (**Figure 2b**). The first five sampling locations are in the central valley and are characterized by vegetation typical of TRC protected slopes and/or bottomland forest, as described above. North Gate has extensive *L. maackii* invasion and is distinct from the other valley locations for its close proximity to the Meramec River. Mincke Quarry has been modified by mining activities and has mostly shallow soils, exposed slopes, *J. virginiana* and *L. maackii* invasion. Mincke Valley contains remnants of old building foundations but is otherwise closed-canopy oak-hickory forest with understory dominated by *A. triloba* and *L. maackii*. Plot 7/8 is on a steep west-facing slope dominated by oak-hickory forest. Ticks were captured using a combination of standard drag sampling, flagging, and dry ice trapping methods, as described in the CDC tick surveillance methods guide [30].

Field collected ticks were transported alive to the TRC laboratory and frozen at −80°C. There, ticks were identified and sorted according to species, sex, and life stage on a cold tray under a dissecting microscope using taxonomic keys [31, 32]. Ticks were then grouped into pools (up to n = 5 for adult ticks and n = 25 for nymphs) by location, sampling date, species, sex and life stage. The pools were stored at −80°C and transported on dry ice to Washington University School of Medicine for further processing.

### Quantification by real-time RT-qPCR

For each RT-qPCR assay, we used *in vitro* transcribed RNAs encompassing the target sequences to define the efficiency and limit of detection of the assay. These targets included sequences specific to BRBV nucleoprotein (BRBV NP) [12], HRTV small segment (HRTV S) [33], and lone star tick 16S mitochondrial ribosomal RNA (rRNA). Standards were diluted to obtain 2.5 × 10^7^ RNA copies/μL and this was 10-fold serially diluted to 0.025 copies/μL. 4 μL of the diluted standards was used to generate standard curves in a quantitative RT-PCR (RT-qPCR) employing the following conditions; 48 °C for 15 min, 95 °C for 10 min, (95 °C for 15 s, 60 °C for 1 min) X 49 cycles on the QuantStudio 6 flex Real-time PCR system (Applied Biosystems), using the TaqMan RNA-to-CT 1-step kit (Applied Biosystems).

### RNA Extraction and Virus Detection

1.0 mL of cold Trizol reagent was added (Invitrogen, cat # 15596018) to a 2 mL homogenization tube containing three stainless steel beads and the pooled ticks. For every five pools, we added a negative control tube (without ticks) as a control for cross contamination. The tubes were then placed in the TissueLyser II (Qiagen Cat. No. / ID: 85300) and run at 30 Hz (1800 oscillations/minute) at 30 secs interval until ticks were well homogenized. The homogenates were then subjected to RNA extraction using the manufacturer’s protocol. The extracted RNA was stored at −80°C until ready to be used. All RNA samples were screened separately in a one-step RT-qPCR for the presence of BRBV and HRTV viral RNA with primers and probes listed in **Table 1**. An experiment (sets of 5 samples plus 1 control) was considered valid if the negative control (samples extracted without ticks) had a Ct value of undetermined, and pools were scored as positive if it had a Ct value equal to or less than 35 (Ct ≤ 35) for both 16S and virus-specific primers.

**Table 1.**
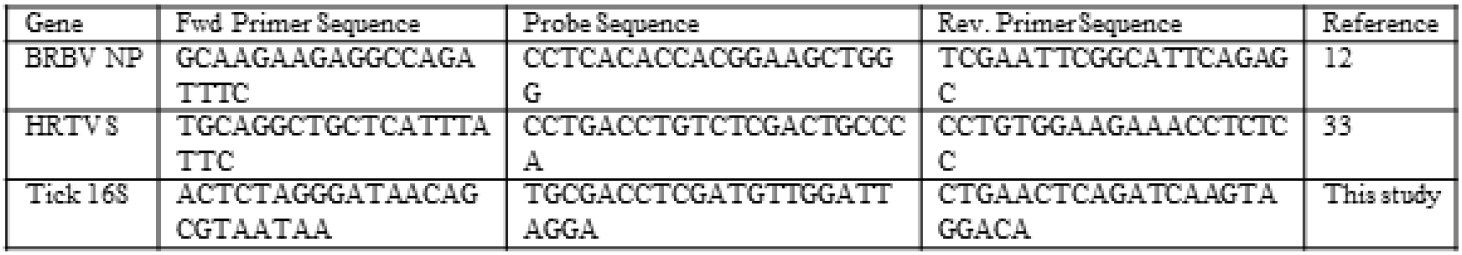
Real time PCR primer-probes for the detection of BRBV HRTV and tick 16S rRNA RNA

### Maximum likelihood Estimate (MLE) of Infection Rate (IR)

Maximum likelihood estimate (MLE) of the infection prevalence per 1,000 ticks at 95% confidence intervals (CIs) for the infection rate were computed using the Excel Add-In [34].

## RESULTS

### Evaluation of RT-qPCR primers and probes for BRBV and HRTV

We used published primers and probes targeting BRBV nucleoprotein (BRBV-NP) [12] and HRTV small segment (HRTV-S) [33] for virus RNA detection. A *de novo A. americanum* tick 16S mitochondrial rRNA (Tick 16S) primer/probe was used to quantify tick RNA, which also served as an indicator for successful tick RNA extraction using our homogenization/extraction method. To define the sensitivity of these primers, we used serial dilutions of the *in vitro* transcribed target RNA sequences standard as template and reliably detected 10 RNA copies in the reaction at an average Ct value of 36 for BRBV-NP and 16S primer/probe sets, and 35 for HRTV-S (**Figure 1a**). In order to mimic physiological conditions and to determine the presence of RT-qPCR inhibitors in the tick homogenates, we spiked RNA extracted from a BRBV/HRTV-negative tick pool with our RNA standard and generated a standard curve as above and still detected 10 RNA copies at an average Ct value of 36 for BRBV-NP and 16S and 35 HRTV-S (**Figure 1b**), suggesting that RNA from homogenized ticks does not inhibit the RT-qPCR reaction. Based on this data, we set the cut off Ct-value for a positive sample at ≤ 35.

**Figure 1:**
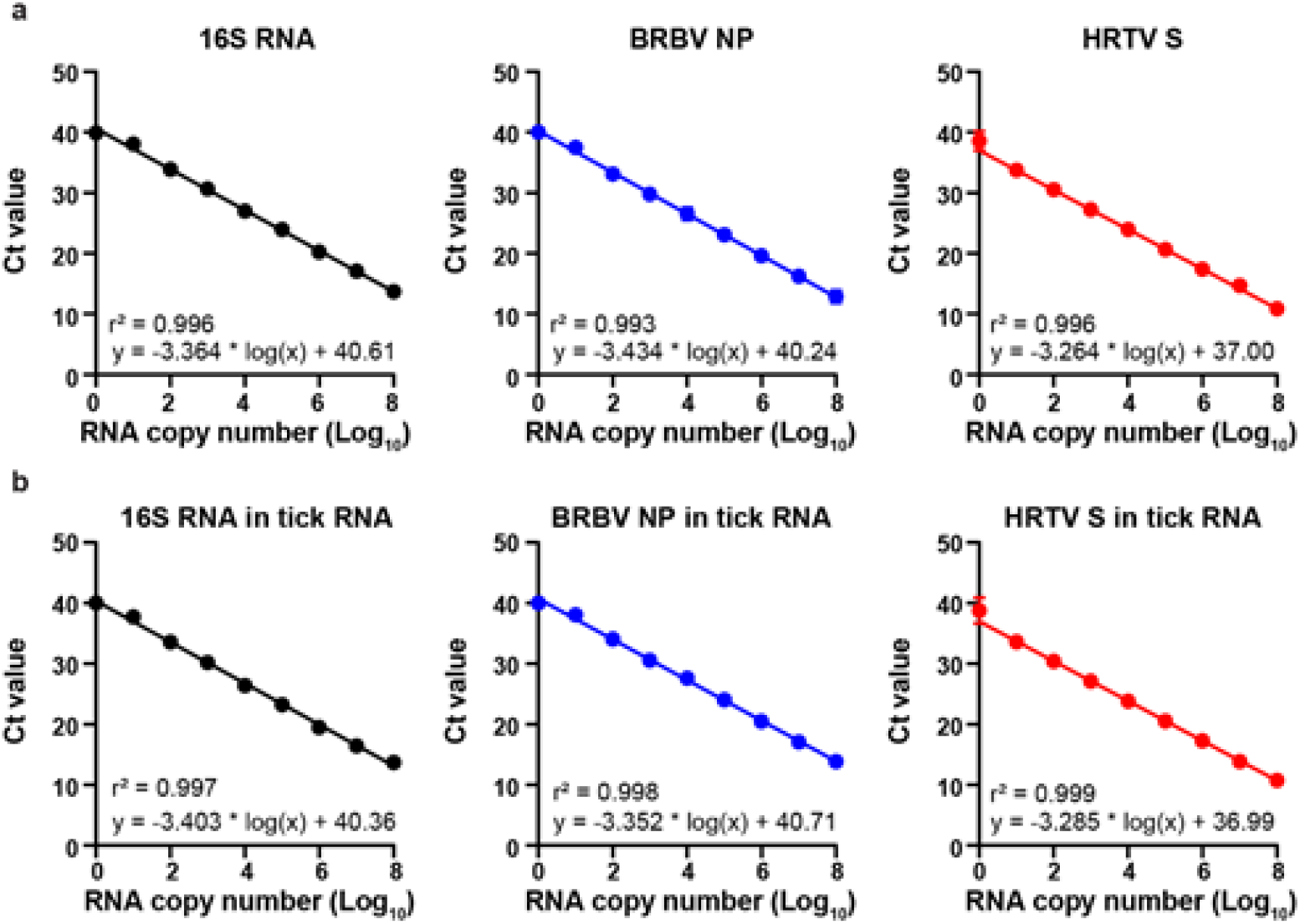
Sensitivity of Primer/Probes in detecting *in vitro* transcribed RNA Standard. The primer/probe sets 16S (*Left*), BRBV-NP (*Middle*) and HRTV-S (*Right*) were used to generate a standard curve in RT-qPCR using a 10 fold serially diluted *in vitro* transcribed RNA as template **(a)** or tick RNA spiked with *in vitro* transcribed RNA **(b)**.

**Figure 2:**
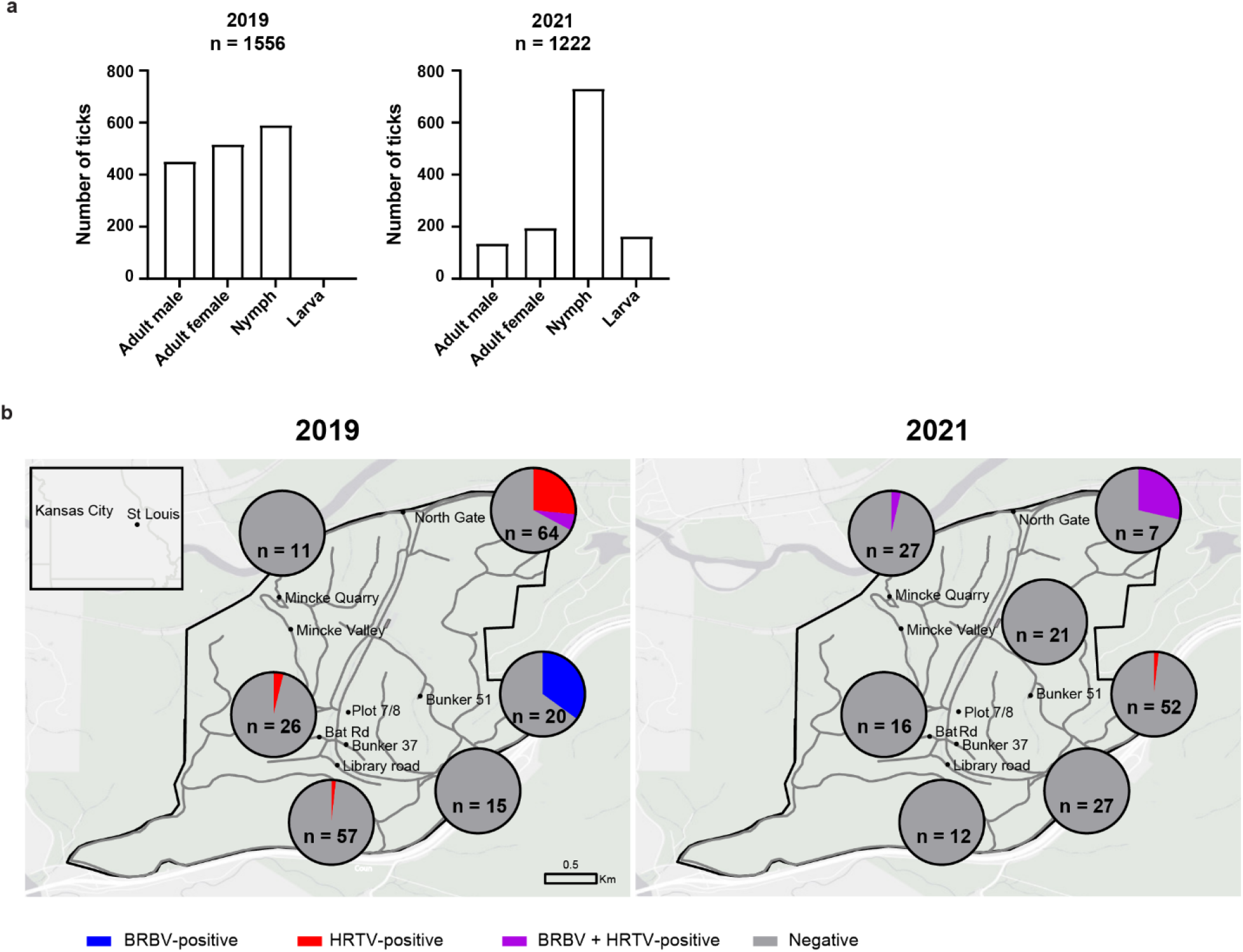
Number of ticks collected in study, grouped according life stage and sex. Number of ticks collected in study, grouped according life stage and sex in 2019) **Figure 2a** (left panel) and 2021 (right panel). **Figure 2b Positivity of BRBV and HRTV by location during 2019 and 2021 collections.** Map of TRC showing 8 approximate tick sampling locations during 2019 and 2021 sampling seasons. In 2019, the locations sampled were Bunker 51, Bat Road, Bunker 37, North Gate, Mincke Quarry and Library Road. In 2021, we sampled all 2019 locations except Bunker 37, and two new locations - Mincke Valley and Plot 7/8. The pie charts represent the number of pools positive for BRBV (blue), HRTV (red), BRBV & HRTV (purple), or negative (grey), and the geographical location of these virus positive pools. The tick pool number is shown inside the pie charts. The black outline shows the TRC property boundary and gray lines depict paved and gravel roads within TRC property. The insert map is that of US State of Missouri with the location of TRC (black dot) relative to Kansas City, and St. Louis city. Maps were created using ArcGIS Pro v2.8 [37] with included area basemap and spatial data collected at TRC.

### Tick collection

Ticks were collected in 2019 (n = 1556) and in 2021 (n = 1222) from eight different locations at TRC. *A. americanum* accounted for nearly 99.4% of the collected ticks at TRC in both 2019 and 2021 collections. The remaining 0.6% were *Dermacentor variabilis*. In the 2019 collection, 29% of *A. americanum* ticks were adult males, 33% were adult females, and 38% were nymphs. In the 2021 collection, 11% were adult males, 16% were adult females, 13% larva, and 60% were nymphs (**Figure 2a**).

### Detection of BRBV and HRTV in Ticks

In 2019, 11 and 23 pools out of the 193 tested were BRBV and HRTV positive respectively. Seven of the BRBV positive pools were from Bunker 51 and ticks were collected on the 23^rd^ of May, and the remaining 4 BRBV positive pools were made up of ticks collected on the 10^th^ of June at North Gate. Of the 11 BRBV positive pools, 8 were pools of adult males, 2 of adult females and 1 of nymphs. Of the 23 HRTV positive pools, 21 were from North Gate and one each from Library Road and Bat Road collected on the 10^th^, 3^rd^ and 4^th^ of June respectively. The HRTV positive pools included 20 pools of adult males, 1 of adult females and 2 of nymphs (**Figure 2b, Table 2**). The infection rates (IRs) for *A. americanum* among the virus positive locations for BRBV were 7.0 (CI = 2.3 - 16.7) for North Gate and 80.6 (CI = 36.5 – 155.4) for Bunker 51. The HRTV IRs were, 39.5 (CI = 26.6 – 57.2) for North Gate, 8.1 (CI = 0.47 – 38.6) for Bat Road, and 2.8 (CI = 0.16 – 13.7) for Library Road. We also calculated IRs of BRBV and HRTV in the ticks by life stage and sex. The BRBV and HRTV IRs for all adults (male and female) *A. americanum* were 10.5 (5.4 – 18.5) and 22.6 (14.6 – 33.6) respectively. The BRBV IRs for *A. americanum* adult male, adult female and nymph in the entire 2019 collection were 18.3 (8.6 – 34.5), 3.8 (0.70 – 12.5) and 1.7 (0.10 – 8.3) respectively. HRTV IRs were 48.8 (31.0 – 73.4), 1.9 (0.1 – 9.2) and 3.4 (0.6 – 11.4) for male, female and nymph respectively.

**Table 2.** Information on the virus positive pools in the 2019 collection

| Location | Male |  |  | Female |  |  | Nymph |  |  | Total |  |
| --- | --- | --- | --- | --- | --- | --- | --- | --- | --- | --- | --- |
|  | HRTV<br>+ | BRBV<br>+ | #pools | HRTV<br>+ | BRBV<br>+ | #pools | HRTV<br>+ | BRBV<br>+ | #pools | #+<br>pools | #<br>pools |
| Bunker 51 | 0 | 6 | 11 | 0 | 1 | 9 | 0 | 0 | 0 | 7 | 20 |
| Bat Road | 1 | 0 | 13 | 0 | 0 | 13 | 0 | 0 | 0 | 1 | 26 |
| Bunker 37 | 0 | 0 | 7 | 0 | 0 | 5 | 0 | 0 | 3 | 0 | 15 |
| North Gate | 19 | 2 | 27 | 1 | 1 | 25 | 1 | 1 | 12 | *21 | 64 |
| Mincke<br>Quarry | 0 | 0 | 6 | 0 | 0 | 1 | 0 | 0 | 4 | 0 | 11 |
| Library<br>Road | 0 | 0 | 27 | 0 | 0 | 26 | 1 | 0 | 4 | 1 | 57 |
| Total pools | 20 | 8 | 91 | 1 | 2 | 79 | 2 | 1 | 23 | 30 | 193 |
| Indv. Ticks | 450 |  |  | 516 |  |  | 590 |  |  | 1556 |  |

\* 4 pools were positive for both Bourbon and Heartland viruses

In 2021, there were four virus positive pools out of a total of 162 tested. One pool was HRTV positive and the remaining three were positive for both viruses. The virus positive pools were from three locations: Bunker 51 (1/4); North Gate (2/4); and Mincke Quarry (1/4) (**Figure 2b & Table 3**) The ticks were collected on 4^th^ of June, 10^th^ of June and 9^th^ of July respectively. The IRs of BRBV in ticks by life stage and sex were 10.3 (1.85 - 33.2) for adult female, 1.3 (0.008 - 6.5) for nymph, and zero for adult male and larvae. The IRs for HRTV in all adults (male and female) *A. americanum* were 6.0 (1.0 – 19.6) and 2.7 (0.5 – 8.9) for nymphs.

**Table 3.**
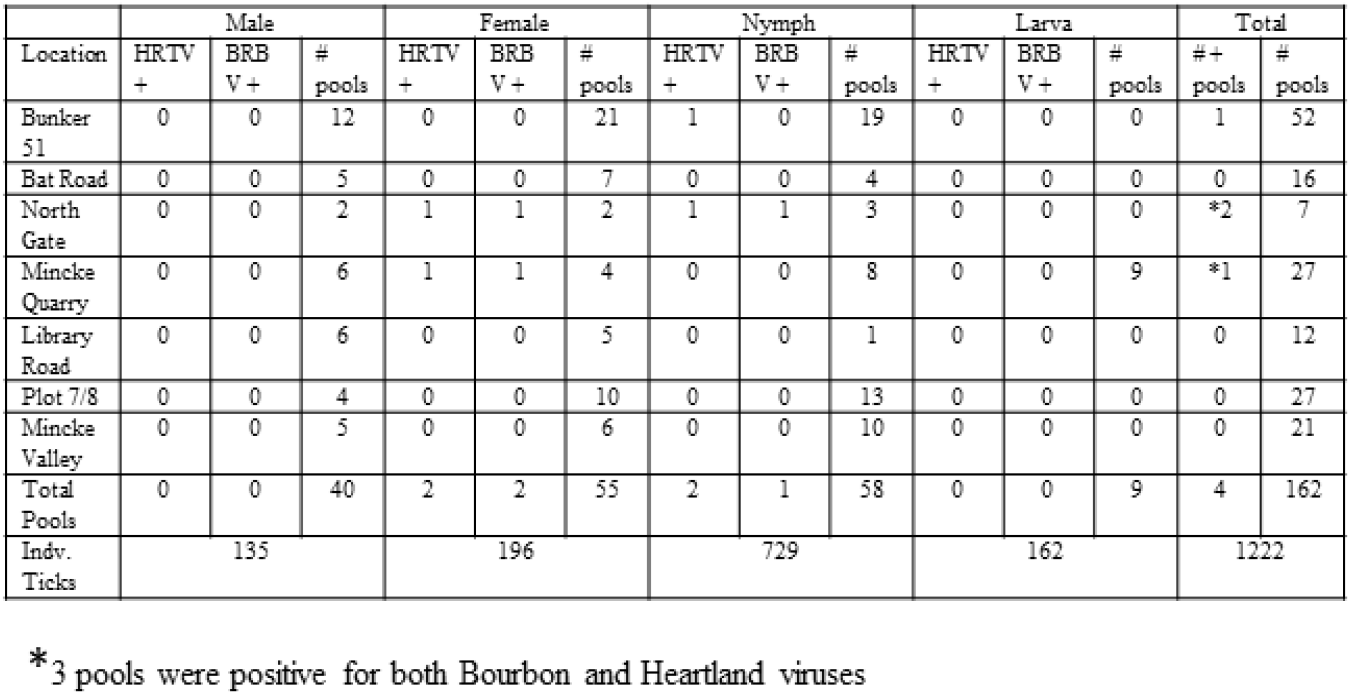
Information on the virus positive pools in the 2021 collection

The BRBV IRs for *A. americanum* adult male, adult female and nymph in the entire study (2019 & 2021) were 14.0 (6.6 – 26.4), 5.6 (1.8 – 13.4) and 1.5 (0.2 – 4.9) respectively and for HRTV for the same period were 36.7 (23.2 – 55.3), 4.2 (1.1 – 11.3) and 3.1 (1.0 – 7.4).

The Ct value for the positive pools ranged from 24.58 - 34.65 corresponding to 6.6 × 10^4^ – 67 RNA copies for BRBV and 23.05 – 35.04 corresponding to 2.0 × 10^4^ – 5.6 RNA copies for HRTV.

## DISCUSSION

In this study we document the presence of BRBV and HRTV viruses in field collected ticks at an environmental field station in the eastern-central part of MO at the edge of the St. Louis metropolitan area. A total of 2778 ticks were collected between May - July of 2019 and 2021, and we detected BRBV and HRTV viral RNA in tick homogenates by RT-qPCR. Thirty-four RT-qPCR virus-positive pools were obtained from five of the eight sampled locations (**Figure 2b**) and were composed of adult (male and female) and nymphal *A. americanum* ticks.

Detection of HRTV and BRBV in adult and nymphal *A. americanum* ticks in our study is consistent with prior studies [14, 15, 18, 26, 33] and support the notion that *A. americanum* is a vector for HRTV and BRBV to humans. The fact that viruses were detected in both nymphal and adult stages exhibiting host-seeking behavior suggest that both life stages could potentially transmit the virus to humans. Host-seeking nymph and adult A*. americanum* ticks, likely acquire the virus during co-feeding with other infected ticks or by feeding on viraemic vertebrate hosts [35, 36]. Transmission could be transstadial, from infected nymphs to adults or from larvae to nymphs, although screening 162 larvae (the life stage with the smallest sample size in our study), did not result in the detection of virus.

BRBV and HRTV were detected at North Gate in adult and nymphal *A. americanum* for both years in our study (**Figure 2b, Tables 2 & 3**) suggesting that these viruses are maintained in the tick and/or host populations in this area of TRC and that an overlap in BRBV and HRTV transmission cycles exist at this location.

The detection of more HRTV positive pools than BRBV observed in this current study was also observed in similar studies in Northwestern Missouri [18, 26, 33] and Kansas [14, 15]. This could be due to BRBV being less efficiently passed between ticks during co-feeding or by transstadial transmission from one life stage to another, or vertically from infected female to offspring, or that BRBV utilizes a vertebrate host(s) that is less abundant or encountered in TRC.

A limitation of our study is the heterogeneity in sampling depth per location. Also because the whole tick was homogenized in Trizol for RNA purification, we could not culture live virus from these homogenates. We were also unsuccessful in amplifying whole gene-segment(s) of BRBV and HRTV, presumably due to the viral RNA abundance.

Additional tick collection efforts combined with animal and human sero-surveillance is important in gaining a deeper understanding of the environmental determinants of vector positivity and risk of exposure of these emerging viruses to humans in the US.

## ACKNOWLEDGEMENTS

We thank the following TRC technicians and undergraduate and high school research fellows for their assistance with tick collection and identification: Althea Bartz, Eleanor Hohenberg, Elise Nishikawa, Rachel Novick, Elora Robeck, Rossana Romo, Laura Tayon, Christopher Tomera, Will Slatin, Angela Yokley. We thank TRC for support and Susan Flowers for running the research fellowship programs. This study was supported with U01-AI151810 and R21-AI151170 grants to ACMB and a Living Earth Collaborative seed grant to SAA, ACMB, and DW.

## REFERENCES

1. Sabin, N.S., et al., Implications of human activities for (re)emerging infectious diseases, including COVID-19. J Physiol Anthropol, 2020. 39(1): p. 29.

2. Jones, K.E., et al., Global trends in emerging infectious diseases. Nature, 2008. 451(7181): p. 990–3.

3. Eisen, R.J., et al., Tick-borne zoonoses in the United States: persistent and emerging threats to human health. ILAR journal, 2017. 58(3): p. 319–335.

4. Rodino, K.G., E.S. Theel, and B.S. Pritt, Tick-Borne Diseases in the United States. Clinical Chemistry, 2020. 66(4): p. 537–548.

5. Rosenberg, R., et al., Vital signs: trends in reported vectorborne disease cases—United States and Territories, 2004–2016. Morbidity and Mortality Weekly Report, 2018. 67(17): p. 496.

6. Rodino, K.G., E.S. Theel, and B.S. Pritt, Tick-Borne Diseases in the United States. Clin Chem, 2020. 66(4): p. 537–548.

7. Tokarz, R. and W.I. Lipkin, Discovery and Surveillance of Tick-Borne Pathogens. Journal of Medical Entomology, 2020. 58(4): p. 1525–1535.

8. Sonenshine, D.E., Range Expansion of Tick Disease Vectors in North America: Implications for Spread of Tick-Borne Disease. Int J Environ Res Public Health, 2018. 15(3).

9. Molaei, G., et al., Bracing for the Worst — Range Expansion of the Lone Star Tick in the Northeastern United States. New England Journal of Medicine, 2019. 381(23): p. 2189–2192.

10. Bosco-Lauth, A.M., et al., Vertebrate Host Susceptibility to Heartland Virus. Emerg Infect Dis, 2016. 22(12): p. 2070–2077.

11. Brault, A.C., et al., Heartland Virus Epidemiology, Vector Association, and Disease Potential. Viruses, 2018. 10(9).

12. Lambert, A.J., et al., Molecular, serological and in vitro culture-based characterization of Bourbon virus, a newly described human pathogen of the genus Thogotovirus. J Clin Virol, 2015. 73: p. 127–132.

13. Newman, B.C., et al., Heartland Virus in Lone Star Ticks, Alabama, USA. Emerg Infect Dis, 2020. 26(8): p. 1954–1956.

14. Savage, H.M., et al., Surveillance for Tick-Borne Viruses Near the Location of a Fatal Human Case of Bourbon Virus (Family Orthomyxoviridae: Genus Thogotovirus) in Eastern Kansas, 2015. J Med Entomol, 2018. 55(3): p. 701–705.

15. Savage, H.M., et al., Surveillance for Heartland and Bourbon Viruses in Eastern Kansas, June 2016. J Med Entomol, 2018. 55(6): p. 1613–1616.

16. Kosoy, O.I., et al., Novel thogotovirus associated with febrile illness and death, United States, 2014. Emerg Infect Dis, 2015. 21(5): p. 760–4.

17. McMullan, L.K., et al., A new phlebovirus associated with severe febrile illness in Missouri. New England Journal of Medicine, 2012. 367(9): p. 834–841.

18. Savage, H.M., et al., Bourbon Virus in Field-Collected Ticks, Missouri, USA. Emerg Infect Dis, 2017. 23(12): p. 2017–2022.

19. Bricker, T.L., et al., Therapeutic efficacy of favipiravir against Bourbon virus in mice. PLoS Pathog, 2019. 15(6): p. e1007790.

20. Dupuis, A.P., 2nd, et al., Heartland Virus Transmission, Suffolk County, New York, USA. Emerg Infect Dis, 2021. 27(12).

21. Tuten, H.C., et al., Heartland Virus in Humans and Ticks, Illinois, USA, 2018-2019. Emerg Infect Dis, 2020. 26(7): p. 1548–1552.

22. Jackson, K.C., et al., Bourbon Virus in Wild and Domestic Animals, Missouri, USA, 2012-2013. Emerg Infect Dis, 2019. 25(9): p. 1752–1753.

23. Butenko, A.M., et al., [Dhori virus--a causative agent of human disease. 5 cases of laboratory infection]. Vopr Virusol, 1987. 32(6): p. 724–9.

24. Frese, M., et al., Human MxA protein inhibits tick-borne Thogoto virus but not Dhori virus. J Virol, 1995. 69(6): p. 3904–9.

25. Tran, N.T.B., et al., Zoonotic Infection with Oz Virus, a Novel Thogotovirus. Emerg Infect Dis, 2022. 28(2): p. 436–439.

26. Savage, H.M., et al., Surveillance for Heartland Virus (Bunyaviridae: Phlebovirus) in Missouri During 2013: First Detection of Virus in Adults of Amblyomma americanum (Acari: Ixodidae). J Med Entomol, 2016. 53(3): p. 607–612.

27. Liu, Q., et al., Severe fever with thrombocytopenia syndrome, an emerging tick-borne zoonosis. Lancet Infect Dis, 2014. 14(8): p. 763–772.

28. Van Horn, T.R., et al., Landscape Physiognomy Influences Abundance of the Lone Star Tick, Amblyomma americanum (Ixodida: Ixodidae), in Ozark Forests. Journal of Medical Entomology, 2018. 55(4): p. 982–988.

29. Kensinger, B.J. and B.F. Allan, Efficacy of dry ice-baited traps for sampling Amblyomma americanum (Acari: Ixodidae) varies with life stage but not habitat. J Med Entomol, 2011. 48(3): p. 708–11.

30. Centers for Disease Control and Prevention. 2020. Guide to the Surveillance of Metastriate Ticks (Acari: Ixodidae) and their Pathogens in the United States. Division of Vector□Borne Diseases, CDC. Atlanta & Ft. Collins. [April 2020; www.cdc.gov/ticks/surveillance].

31. Keirans, J.E. and L.A. Durden, Illustrated key to nymphs of the tick genus Amblyomma (Acari: Ixodidae) found in the United States. J Med Entomol, 1998. 35(4): p. 489–95.

32. Keirans, J.E. and T.R. Litwak, Pictorial key to the adults of hard ticks, family Ixodidae (Ixodida: Ixodoidea), east of the Mississippi River. J Med Entomol, 1989. 26(5): p. 435–48.

33. Savage, H.M., et al., First detection of heartland virus (Bunyaviridae: Phlebovirus) from field collected arthropods. Am J Trop Med Hyg, 2013. 89(3): p. 445–452.

34. Biggerstaff, B., PooledInfRate, Version 3.0: a Microsoft Excel Add-In to compute prevalence estimates from pooled samples. Centers for Disease Control and Prevention, Fort Collins, CO, 2006.

35. Godsey, M.S., et al., Experimental Infection of Amblyomma americanum (Acari: Ixodidae) With Bourbon Virus (Orthomyxoviridae: Thogotovirus). J Med Entomol, 2021. 58(2): p. 873–879.

36. Godsey, M.S., et al., Transmission of Heartland Virus (Bunyaviridae: Phlebovirus) by Experimentally Infected Amblyomma americanum (Acari: Ixodidae). J Med Entomol, 2016. 53(5): p. 1226–1233.

37. Citation: ESRI 2021. ArcGIS Pro version 2.8. Redlands, CA: Environmental Systems Research Institute.

